# Viral apoptosis evasion via the MAPK pathway by use of a host long noncoding RNA

**DOI:** 10.1101/153130

**Authors:** Samantha Barichievy, Jerolen Naidoo, Mikaël Boullé, Janine Scholefield, Suraj P. Parihar, Anna K. Coussens, Frank Brombacher, Alex Sigal, Musa M. Mhlanga

## Abstract

An emerging realisation of infectious disease is the high incidence of genetic instability resulting from pathogen-induced DNA lesions, often leading to classical hallmarks of cancer such as evasion of apoptosis. The Human Immunodeficiency Virus type 1 (HIV-1) induces apoptosis in CD4+ T cells but is largely non-cytopathic in macrophages, thereby leading to long-term dissemination of the pathogen specifically by these host cells. Apoptosis is triggered by double-strand breaks (DSBs), such as those induced by integrating retroviruses, and is coordinated by the p53-regulated long noncoding RNA lincRNA-p21, in a complex with its protein binding partners HuR and hnRNP-K. Here, we monitor the cellular response to infection to determine how HIV-1 induces DSBs in macrophages yet evades apoptosis in these cells. We show that the virus does so by securing the pro-survival MAP2K1/ERK2 cascade early upon entry, in a gp120-dependent manner, to orchestrate a complex dysregulation of lincRNA-p21. By sequestering HuR in the nucleus, HIV-1 enables lincRNA-p21 degradation. Simultaneously, the virus permits transcription of pro-survival genes by sequestering hnRNP-K in the cytoplasm via the MAP2K1/ERK2 pathway. Notably, this pro-survival cascade is unavailable for similar viral manipulation in CD4+ T cells. The introduction of MAP2K1, ERK2 or HDM2 inhibitors in HIV-infected macrophages results in apoptosis providing strong evidence that the viral-mediated apoptotic block can be released, specifically by restoring the nuclear interaction of lincRNA-p21 and hnRNP-K. These results reveal pathogenic control of apoptosis and DNA damage via a host long noncoding RNA, and present MAP2K1/ERK2 inhibitors as a novel therapeutic intervention strategy for HIV-1 infection in macrophages.

## INTRODUCTION

Microbial infection, particularly in the case of viral pathogens, often leads to genetic changes in the host chromatin, either via an increase in DNA damage or a decrease in DNA repair activity (Weitzman and Weitzman, 2014). Such genetic instabilities can develop into infection-induced malignancies (de Martel et al., 2012), and are repeatedly linked to classic cancer hallmarks such as enhanced cellular proliferation and evasion of apoptosis (Hanahan and Weinberg, 2011). For example, the E7 protein from the Human Papillomavirus (HPV) directly binds the Retinoblastoma tumour suppressor protein leading to a downstream block in p53-dependent apoptosis (Moody and Laimins, 2010). Similarly, the EBNA3C protein from Epstein Barr Virus (EBV) directly enhances Aurora kinase B activity resulting in aberrant cell division and downregulation of apoptosis in B cells, despite accumulating DNA damage (Jha et al., 2013).

In all cases observed thus far, DNA damage initiated by pathogenic infection triggers damage sensor proteins that launch a cascade to recruit repair proteins and form repair response foci (Ciccia and Elledge, 2010; Jackson and Bartek, 2009). This is particularly important in response to double-strand breaks (DSBs), which are the most detrimental form of DNA lesion leading to apoptosis (Jackson and Bartek, 2009). The MRE11/RAD50/NBS1 (MRN) complex senses DSBs leading to activation of the ataxia telangiectasia mutated (ATM) protein kinase and the tumour suppressor protein p53 (Jackson and Bartek, 2009; Meek, 2004).

Various viruses induce DSBs following infection, and they have evolved a plethora of mechanisms to counteract the ensuing host innate immune response (Weitzman and Weitzman, 2014). This includes viruses that are not directly implicated in causing cancer, although they promote the accumulation of genetic lesions, and target oncogenic hallmark pathways such as apoptosis (Niller et al., 2011). HIV-1 is one such virus, which causes apoptosis in CD4+ T cells driven by the induction of DSBs during viral-mediated integration (Cooper et al., 2013; Doitsch et al., 2010). The progressive loss of CD4+ T cells is so reliable that it is used as a marker of HIV-1 disease progression. Intriguingly, macrophages are also permissive to HIV-1 infection but are largely spared the cytopathic effects of replicating virus, suggesting a selective impairment of the apoptotic response in these specific host cells (Cummins and Badley, 2013). Several viruses, including HIV-1, have been shown to prevent TRAIL-induced apoptosis in macrophages via the control of certain host cytokines (Swingler et al., 2007). However, the roles of cellular integration-associated DSBs and p53 as well as the fundamental molecular mechanism underlying evasion of apoptosis remains unknown.

It is now understood that p53 outsources a critical portion of the apoptotic response to the long noncoding RNA, lincRNA-p21 (Huarte et al., 2010). In a nuclear complex with its protein binding partner hnRNP-K, lincRNA-p21 orchestrates the apoptotic trigger by specifically repressing the transcription of pro-survival p53 target genes in the nucleus (Huarte et al., 2010). One important target of this complex is MAP2K1 gene expression, whose protein product specifically phosphorylates ERK2 to maintain a cellular survival cascade (Chang and Karin, 2001). Interestingly, activated ERK2 also associates with the HIV-1 pre-integration complex (PIC) thus facilitating successful integration in macrophages (Jacque et al., 1998; Bukong et al., 2010). In contrast, JNK and Pin1 (and importantly not ERK2) facilitate HIV-1 integration in activated CD4+ T cells (Manganaro et al., 2010), possibly because ERK2 expression is specifically and selectively shut down as dual positive (CD4+CD8+) cells go through selection and become CD4+ T cells (Fischer et al., 2005; Chang et al., 2012). This renders ERK2 unavailable for viral use in CD4+ T cells. Clearly, for HIV-1 to evade apoptosis following integration while simultaneously ensuring cellular survival, the virus would need to carefully manipulate these pathways. Both p53-mediated apoptosis, and ERK2-mediated cellular survival, share an intimate connection at the intersection of lincRNA-p21 and MAP2K1. This junction also has a critical connection to HIV via the MAP2K1/ERK2 requirement for viral integration in macrophages. Thus HIV-1 must have evolved a pivotal mechanism to control these host factors as a means to evade apoptosis, with lincRNA-p21 as a central target.

Here we show how during viral infection of macrophages, HIV-1 is able to cause DSBs without the infected cells apoptosing. We dissect the mechanism of apoptosis evasion and show that HIV-1, in a gp120-dependent manner, triggers sequestration of specific protein binding partners of lincRNA-p21 to different cellular compartments. This ensures lincRNA-p21 transcripts are continually degraded, while simultaneously preventing formation of an active pro-apoptotic nuclear complex. By using a variety of interventions, including anticancer therapeutics, we were able to reverse these effects and demonstrate how control of lincRNA-p21 activity is critical to the apoptosis evasion process of HIV-1 in macrophages. Given that pathogen-induced genetic instability is fairly common, this mechanism is quite likely to be broadly conserved across several host-pathogen interactions, particularly in the context of infection-induced malignancies.

## RESULTS

### HIV masks DNA damage

As HIV-1 is able to extend survival and prevent elimination of infected macrophages (Swingler, 2007), it is likely that such cells can be exposed to, yet withstand, significant genotoxic stress. Indeed, chronically infected U937 monocyte-like cells are more resistant to DNA-damaging agents (Tanaka et al., 1999). Thus we reasoned that in HIV-infected macrophages, only a portion of the cellular DSB surveillance mechanism may be intact. To ascertain this, we added a lethal dose of Doxorubicin to HIV-infected Ghost(3) reporter cells (possessing an integrated HIV LTR-driven GFP reporter, that allow for real-time monitoring of major infection events, Figures S1A, S1B) or primary monocyte-derived macrophages (Mφ), and assessed the presence of DSBs as well as key elements of the subsequent cellular response pathway. Active viral replication following integration and the related DSBs were visualised in Ghost(3) cells by immunofluorescent staining of H2A.X (Figures 1A, S1C-S1E). This histone variant is rapidly phosphorylated at serine 139 (Ser139) within seconds of such DNA damage and persists until the DSB is repaired (Keogh et al., 2006). By concurrently assessing the production of activated caspase 3 as a marker of apoptosis induction (shown as percentage cell survival), we observed that cells, which received lethal doses of Doxorubicin prior to HIV-1 infection (Doxo. 1st HIV 2nd), underwent apoptosis (~0% survival, Figure 1A lower panel). However, cells infected 24 hours before being exposed to the same duration and concentration of Doxorubicin treatment (HIV 1st Doxo. 2nd) were protected from apoptosis (60% survival, Figure 1A upper panel). Notably, survival in HIV-infected cells exposed to Doxorubicin was the same as HIV-1 infection alone (Figure 1A upper panel), although viral replication was lower in the former conditions (Figure 1B, 16 000 Gag units vs Figure S1B, 48 000 Gag units). Remarkably, the same apoptosis-resistance phenotype was observed in Doxorubicin-treated Mφ across 3 separate donors (Figures 1C) infected at physiologically relevant levels (Figure 1D). These striking data suggested that the virus is able to decouple the DNA damage surveillance mechanism from the apoptotic response in infected cells. In addition the protection from apoptosis afforded by infection does indeed persist despite additional, otherwise lethal challenges to the genomic integrity of these cells.

**Figure 1.**
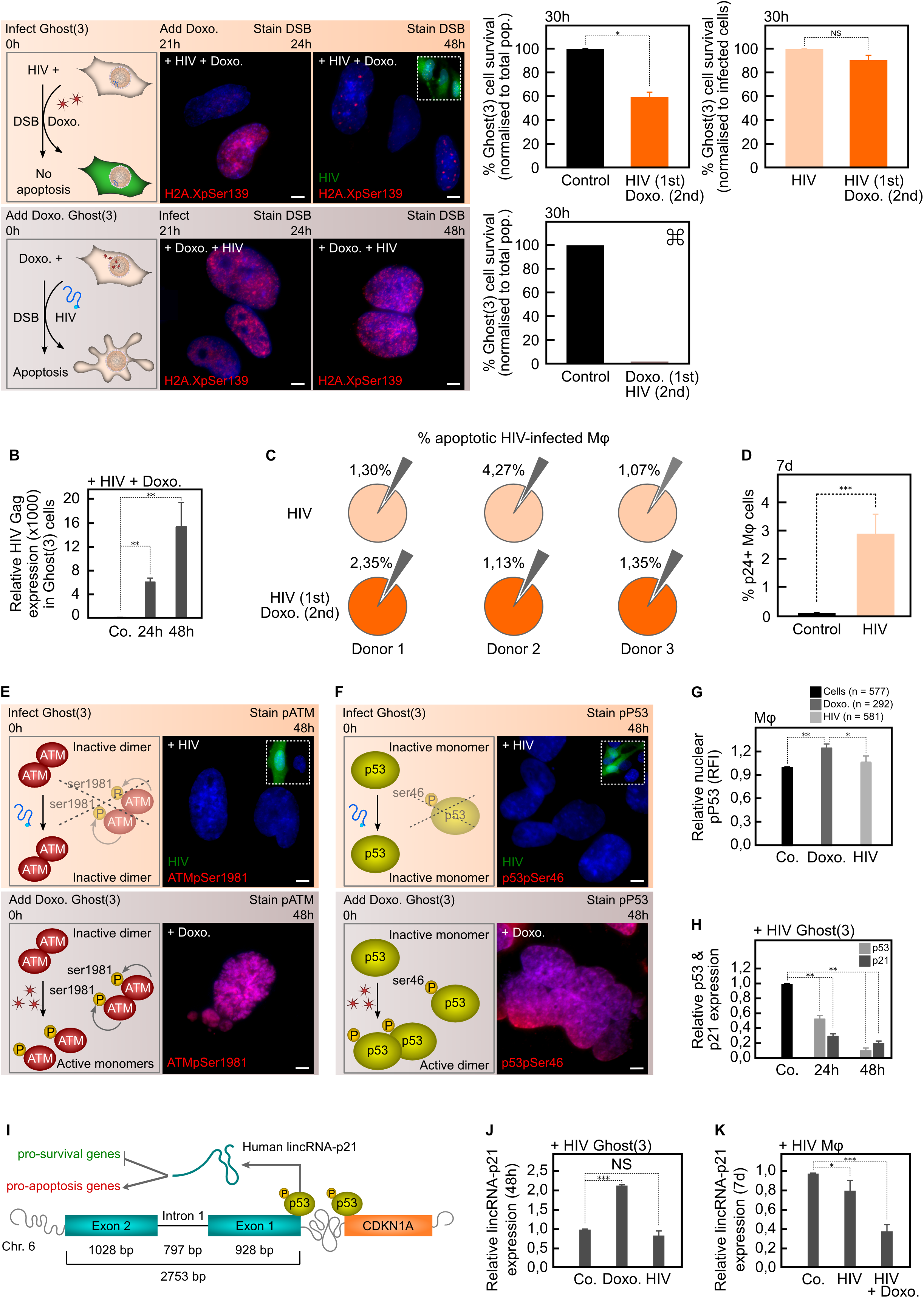
HIV-1 masks DNA damage, protects against additional lethal DNA damage, and prevents lincRNA-p21 upregulation. (A) Ghost(3) cells infected 24 hours prior to Doxorubicin treatment have numerous DSBs as detected by H2A.XpSer139 immunofluorescence staining (+HIV+Doxo.) but do not undergo apoptosis when normalised to infected cells (HIV 1st Doxo. 2nd; orange bars). Cells infected after exposure to Doxorubicin (+Doxo.+HIV) have extensive H2A.XpSer139 staining and do undergo apoptosis. ⌘ Too few attached cells present for statistical analysis (<20). (B) Ghost(3) cells support HIV-1 replication following additional lethal DNA damage (+HIV+Doxo.) as detected by quantitative real-time RT-PCR analysis of HIV-1 Gag expression relative to the HPRT housekeeping gene and normalised to uninfected cells (mean ± SE of 3 biological replicates in triplicate). (C) The percentage of apoptotic HIV-infected Mφ (small grey wedges, upper row) does not significantly increase in HIV-infected Mφ exposed to Doxorubicin (small grey wedges, lower row) across 3 separate donors (biological replicates). (D) Mφ were infected with HIV-1 at a physiologically relevant rate as measured at 7 days by p24 flow cytometry analysis (mean ± SE of 3 biological replicates). (E) Inactive ATM dimers do not undergo autophosphorylation of serine residue 1981 in response to HIV-induced DSBs as measured by immunofluorescence staining (ATMpSer1981) in Ghost(3) cells. Phosphorylated ATM is detected in Doxorubicin-treated cells. (F) Inactive p53 monomers are not phosphorylated at serine residue 46 (specific apoptotic mark) in response to HIV-1 infection of Ghost(3) cells as measured by immunofluorescence staining (p53pSer46). Activated p53 dimers are detected in Doxorubicin-treated cells. (G) The relative nuclear fluorescence intensities (RFI) of p53ser46 residues (specific apoptotic mark) is not increased in response to HIV-1 infection of Mφ, but is increased in Doxorubicin-treated Mφ (mean ± SE of 3 biological replicates). (H) HIV-1 infection significantly decreases p53 and CDKN1A expression over time in Ghost(3) cells as detected by quantitative real-time RT-PCR analysis relative to the HPRT housekeeping gene and normalised to uninfected cells (mean ± SE of 3 biological replicates in triplicate). (I) Human lincRNA-p21, which is located upstream of CDKN1A/p21 (orange) on chromosome 6 and comprised of 2 exons (blue) and a single intron, is transcribed by p53 (yellow) in response to DNA damage. LincRNA-p21 mediates cellular apoptosis by down-regulating pro-survival genes (green) and up-regulating pro-apoptosis genes (red). (J) HIV-induced DNA damage does not lead to enhanced lincRNA-p21 expression in Ghost(3) cells as detected at 48 hours by quantitative real-time RT-PCR analysis relative to the HPRT housekeeping gene and normalised to untreated cells. Doxorubicin treatment does lead to enhanced lincRNA-p21 expression (mean ± SE of 3 biological replicates in triplicate). (K) LincRNA-p21 expression is significantly decreased in HIV-infected Mφ (7 days), with an even greater decrease observed in infected Mφ exposed to Doxorubicin (HIV+Doxo. at 8 days) as detected by quantitative real-time RT-PCR analysis relative to the HPRT housekeeping gene and normalised to untreated cells (mean ± SE of 3 biological replicates in triplicate). Cells were counterstained with DAPI; scale bars = 10μM; two-tailed paired Student T-test, ***p < 0.001, **p < 0.01, *p < 0.05, NS (not significant); see also supplemental figure 1.

The cellular response to DSBs is exquisitely sensitive and DNA damage is recognized by ATM, which is activated by auto-phosphorylation of Ser1981 (Lee and Paull, 2005). A key downstream target of activated ATM is p53, which is phosphorylated at multiple residues (Riley et al., 2008) including Ser46. Phosphorylation at Ser46 specifically directs cellular responses towards apoptosis (Smeenk et al., 2011). As DSBs are known to trigger ATM and p53 activation, we assessed their specific phosphorylation states in response to viral and chemical agent-induced DNA damage. Doxorubicin treatment resulted in the robust phosphorylation of both ATMpSer1981 (Figure 1E) and p53pSer46 (Figure 1F) within 8 hours in the nuclei of Ghost(3). In contrast, HIV-1 infection did not lead to phosphoryation of either of these residues over 48 hours. Similarly, the same major apoptotic mark on nuclear-localised p53 was strikingly enhanced in Doxorubicin-treated Mφ as opposed to those exposed to HIV-1 (Figures 1G, S1F). Furthermore, quantitative real-time RT-PCR analysis revealed that expression of p53, as well as one of its major downstream cell cycle factors CDKN1A/p21 (Wu et al., 2010), was significantly decreased during the same time course in Ghost(3) cells in the presence of virus (Figures 1H).

### HIV negatively affects a host long noncoding RNA regulating apoptosis

In response to DNA damage, p53 transcriptionally activates lincRNA-p21 which is located on chromosome 6 upstream of CDKN1A/p21 in human cells (Figure 1I; Huarte et al., 2010; Yoon et al., 2012). Since HIV-1 had neutralised the apoptotic response, we examined the status of lincRNA-p21 in HIV-infected and control cells. As previously noted (Huarte et al., 2010; Yoon et al., 2012), lincRNA-p21 was enhanced in Doxorubicin-treated cells, but HIV-induced DNA damaged Ghost(3) cells showed no increase in lincRNA-p21 expression (Figure 1J). This was also true of viral-infected Mφ, where even in the presence of additional lethal DNA damage (Doxorubicin treatment), lincRNA-p21 levels were not increased but further decreased (Figure 1K). Overall, these data strongly suggested that HIV-1 may prevent apoptosis by direct or indirect regulation of lincRNA-p21.

Under normal conditions, lincRNA-p21 is rendered unstable through the action of HuR/ELAV1 in the nucleus (Figure 2A; Yoon et al., 2012). This occurs via HuR/ELAV1 mediated degradation of lincRNA-p21 through the recruitment of the Ago2 protein and let-7a mirRNA. We hypothesised that HIV-1 may co-opt this mechanism to maintain lincRNA-p21 instability and thus evade apoptosis. Our hypothesis was initially bolstered by the observation of a significant difference in the cellular location of HuR in Mφ and Ghost(3) cells infected with HIV-1 as compared to chemically induced DSBs (Figures 2B-2C, S2A). Indeed during HIV-1 integration and concomitant induction of DSBs (Figures S1C), HuR remained in the nucleus over the course of infection, presumably maintaining its destabilizing effect on lincRNA-p21. Contrastingly, in Doxorubicin-treated cells, HuR moved from the nucleus into the cytoplasm within 8 hours of treatment, releasing the destabilatory effect on lincRNA-p21 and consistent with our observed increase in lincRNA-p21 expression (Figures 1J-1K).

**Figure 2.**
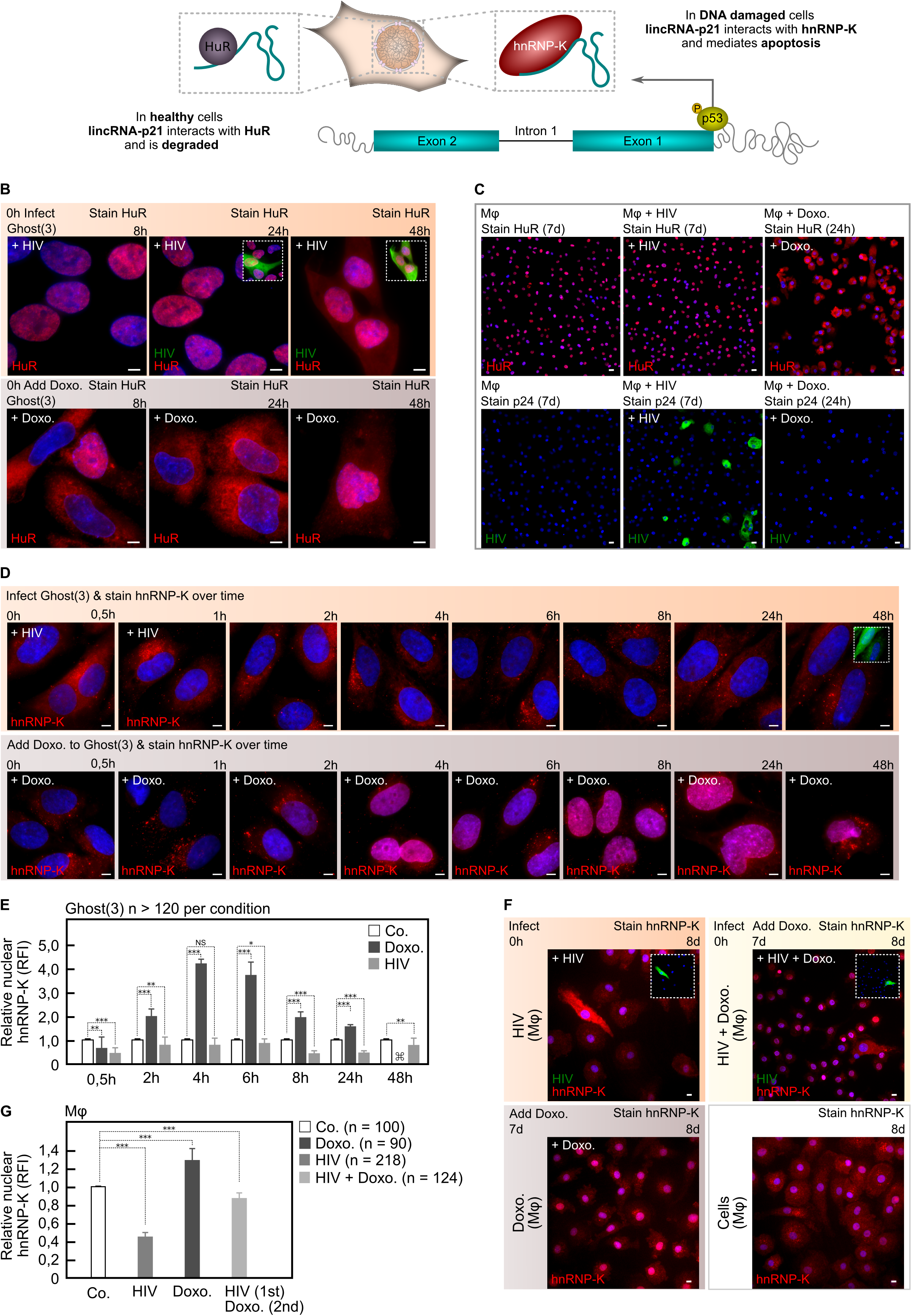
HIV-1 manipulates lincRNA-p21’s protein binding partners. (A) In healthy cells, lincRNA-p21 associates with HuR in the nucleus and is degraded. In response to p53-mediated transcription, lincRNA-p21 associates with hnRNP-K in the nucleus of DNA-damaged cells and regulates apoptosis by localising hnRNP-K to the promoters of p53-repressed genes. (B) Immunofluorescence staining reveals cytoplasmic HuR within 8 hours of Doxorubicin-treated Ghost(3) cells while HIV-infected cells show nuclear HuR up until 48 hours. (C) Immunofluorescence staining reveals nuclear HuR in untreated and HIV-infected Mφ (counterstained with p24 in lower panel), but clear cytoplasmic HuR in Doxorubicin-treated Mφ. (D) Immunofluorescence staining reveals nuclear hnRNP-K within 2 hours of Doxorubicin treatment in Ghost(3) cells, while HIV-infected cells show cytoplasmic hnRNP-K throughout all 8 time points spanning the same 48 hour time course. (E) Quantification of nuclear localised hnRNP-K in HIV-infected Ghost(3) cells over time (mean relative fluorescence intensity (RFI) ± SE of 3 biological replicates). ⌘ Too few attached cells were present for statistical analysis (<20). (F) Immunofluorescence staining reveals nuclear hnRNP-K within 24 hours of Doxorubicin treatment of Mφ (Doxo.), while HIV-infected cells (+HIV) and those subsequently exposed to Doxorubicin (+HIV+Doxo.) show cytoplasmic hnRNP-K. (G) Quantification of nuclear localised hnRNP-K in Mφ infected with HIV-1 or infected and exposed to Doxorubicin (HIV 1st, Doxo. 2nd); mean relative fluorescence intensity (RFI) ± SE of 3 biological replicates). Cells were counterstained with DAPI; scale bars = 10μM; two-tailed paired Student T-test (Figure 2G used two-tailed unpaired Student T-test), ***p < 0.001, **p < 0.01, *p < 0.05, NS (not significant); see also supplemental figure 2.

Since HIV-infected cells failed to apoptose despite lethal levels of DNA damaging agents, and lincRNA-p21 levels were lowered by nuclear resident HuR, we reasoned that silencing HuR should elevate lincRNA-p21 and trigger apoptosis (Yoon et al., 2012). We delivered siRNAs targeted to HuR in HIV-infected Ghost(3) cells and observed a rise in lincRNA-p21 as detected by qRT-PCR (Figures S2B, S2C). However, siHuR-treated cells continued to support HIV-1 replication to the same extent as untreated cells (Figures S2D, S2E). These data suggested that although the virus manipulates HuR location in order to control lincRNA-p21 expression, this alone could not account for the block in apoptosis observed in infected cells. In support of this, exogenously overexpressed full-length lincRNA-p21 was decreased in HIV-infected Ghost(3) cells (Figures S2F, S2G) and did not lead to apoptosis as observed for similarly treated Doxorubicin-induced DNA damaged cells (Figure S2H). Collectively these data pointed to additional molecular mechanisms by which HIV-1 is able to evade apoptosis.

### HIV alters the protein binding partners of a pro-apoptotic host lncRNA

The interplay between p53, lincRNA-p21 and hnRNP-K ensures repression of a specific set of genes that are part of the p53 DNA damage response. Key to this repression is the formation of a lincRNA-p21/hnRNP-K complex in the nucleus (Huarte et al., 2010). Since nuclear localisation of hnRNP-K is required for its repressive function, and as HIV-1 is able to avoid apoptosis, we hypothesised that in addition to reducing nuclear levels of lincRNA-p21, the virus was altering hnRNP-K’s location within infected cells. We monitored hnRNP-K at several time points following DNA damage induced either by Doxorubicin treatment or HIV-1 infection using immunofluorescence staining (Figures 2D-2G). Notably, hnRNP-K was detected in the nuclei of Doxorubicin-treated Ghost(3) cells between 2 and 4 hours post-treatment and by 24 hours, cells were apoptotic. Remarkably, hnRNP-K remained cytoplasmic in HIV-infected cells throughout the duration of the time course with no apoptosis occurring (Figures 2D-2E).

HIV-infected Mφ showed an identical phenotype and furthermore, infected cells subsequently exposed to lethal Doxorubicin treatment showed significantly less nuclear hnRNP-K as compared to untreated cells or those treated with Doxorubicin alone (Figures 2F-2G). These observations parsimoniously explained the inability to trigger transcription of pro-apoptotic genes, since hnRNP-K was unable to associate with lincRNA-p21 in the nucleus. It also explained why the addition of exogenous lincRNA-p21 is unable to induce apoptosis in HIV-infected cells. Indeed, in addition to HuR and hnRNP-K, HIV-1 infection altered the binding of multiple additional protein partners of lincRNA-p21 (assessed by IP-MS, Figures S2I, S2J), lending further support to our hypothesis that HIV-1 alters the lincRNA-p21/hnRNP-K interaction to avert apoptosis.

### HIV controls a pro-apoptotic host lncRNA via MAPK pathway members

MAP2K1 is the primary kinase that activates ERK2 as part of a survival cascade in healthy cells (Habelhah et al., 2001; Cagnol and Chambard, 2010). Following activation, ERK2 specifically phosphorylates hnRNP-K at serines 284 and 353, leading to cytoplasmic accumulation of hnRNP-K (Habelhah et al., 2001). Inhibition of the MAP2K1/ERK2 pathway prevents this phosphorylation of hnRNP-K and thus affects hnRNP-K cellular location (Figure 3A, Habelhah et al., 2001). As MAP2K1 transcription is negatively regulated by the lincRNA-p21/hnRNP-K complex as part of the pro-apoptosis pathway (Huarte et al., 2010), we sought to determine if HIV-1 secures cellular survival via MAP2K1/ERK2. Initially we examined MAP2K1 transcription and found its expression to be significantly up-regulated in the presence of HIV-1 (Figures 3B, S3A). Intriguingly, this was also observed in HIV-infected cells that were subsequently exposed to a lethal dose of Doxorubicin (Figures 3B, S3A). When we added a MAP2K1-specific inhibitor (thus preventing specific downstream ERK2 activation) to HIV-infected cells, MAP2K1 expression was significantly reduced (Figures 3B, S3A). In contrast, infected cells treated with an ERK2-specific inhibitor showed enhanced MAP2K1 expression relative to control cells (Figures 3B, S3A). Together, these observations revealed that HIV-1 infection specifically increases MAP2K1 expression, and strongly suggested that HIV-1 can counteract lincRNA-p21 mediated suppression of MAP2K1. Indeed, when we examined lincRNA-p21 expression under similar conditions, we observed that HIV-infected cells treated with a MAP2K1 inhibitor (hence no activated ERK2) showed significantly increased lincRNA-p21 levels (Figures 3C, S3B). In contrast, infected cells treated with an ERK2 inhibitor did not show increased lincRNA-p21 (Figures 3C, S3B). Notably, in HIV-infected cells subsequently treated with Doxorubicin, lincRNA-p21 levels were also significantly reduced (Figures 3C, S3B). These results were in line with our observed changes in MAP2K1 expression and clearly demonstrated that HIV-1 is able to suppress lincRNA-p21 expression by ensuring high levels of MAP2K1 expression.

**Figure 3.**
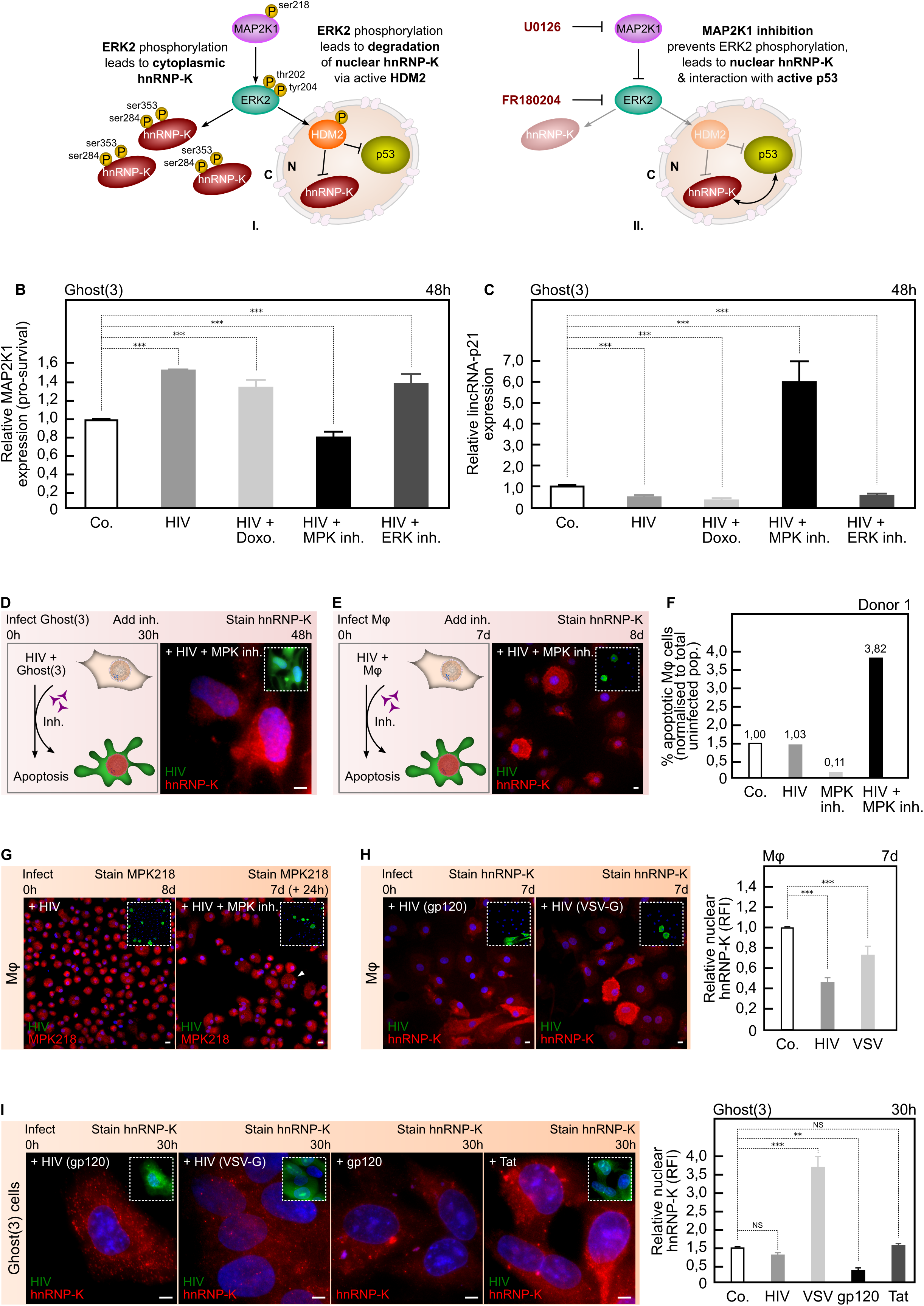
HIV-1 requires MAP2K1/ERK2 and gp120 to ensure hnRNP-K’s cytoplasmic localisation. (A) Activated MAP2K1 specifically phosphorylates ERK2 leading to cytoplasmic accumulation of phosphorylated hnRNP-K (I). Simultaneously, ERK2 activation of HDM2 ensures ubiquitin-mediated degradation of nuclear hnRNP-K and p53 (I). Inhibition of MAP2K1 (or ERK2) prevents ERK2-mediated accumulation of hnRNP-K, and releases the negative regulation of HDM2 on nuclear hnRNP-K and p53 (II). (B) HIV-1 enhances MAP2K1 expression in Ghost(3) cells, in the presence of Doxorubicin (HIV+Doxo.) or an ERK2 inhibitor (HIV+ERK inh.) but not in the presence of a MAP2K1 inhibitor (MPK inh.) as detected by quantitative real-time RT-PCR analysis relative to the HPRT housekeeping gene and normalised to untreated cells (mean ± SE of 3 biological replicates in triplicate). (C) HIV-1 requires enhanced MAP2K1 expression to reduce lincRNA-p21 levels in Ghost(3) cells as only in the presence of a MAP2K1 inhibitor (HIV+MPK inh.) is lincRNA-p21 expression enhanced, as detected by quantitative real-time RT-PCR analysis relative to the HPRT housekeeping gene and normalised to untreated cells (mean ± SE of 3 biological replicates in triplicate). (D) Inhibition of MAP2K1 allows for nuclear localisation of hnRNP-K and apoptosis in HIV-infected Ghost(3) cells as measured by immunofluorescence staining. (E) Inhibition of MAP2K1 allows for nuclear localisation of hnRNP-K and apoptosis in HIV-infected Mφ as measured by immunofluorescence staining. (F) The percentage of apoptotic Mφ (normalised to total uninfected population) increases in HIV-infected Mφ exposed to a MAP2K1 inhibitor (HIV+MPK inh.). Data shown for 1 of 3 donors. (G) Immunofluorescence staining of the Ser218/222 activation mark on MAP2K1 in HIV-infected Mφ shows punctate cytoplasmic localisation of the protein, only in the presence of the inhibitor (white arrowhead). (H) HIV-1 clone NL4-3 packaged with gp120 Env (+HIV gp120) excludes hnRNP-K from the nucleus of infected Mφ as shown by immunofluorescence staining, while HIV-1 packaged with VSV-G (+HIV VSV-G) fails to do so. Quantification of nuclear localised hnRNP-K in infected Mφ shown in right panel (mean relative fluorescence intensity (RFI) ± SE of 3 biological replicates). (I) HIV-1 clone BaL.01 packaged with gp120 Env (+HIV gp120) excludes hnRNP-K from the nucleus of infected Ghost(3) cells as shown by immunofluorescence staining, while HIV-1 packaged with VSV-G (+HIV VSV-G) fails to do so. Cells transfected with a plasmid expressing only HIV-1 gp120 (+gp120) show cytoplasmic hnRNP-K, while control cells transfected with a plasmid expressing only HIV-1 Tat (+Tat) show cytoplasmic and nuclear hnRNP-K. Quantification of nuclear localised hnRNP-K in treated Ghost(3) cells shown in right panel (mean relative fluorescence intensity (RFI) ± SE of 3 biological replicates). Cells were counterstained with DAPI; scale bars = 10μM; two-tailed paired Student T-test, ***p < 0.001, **p < 0.01, *p < 0.05, NS (not significant); see also supplemental figure 3.

We next examined the effects of HIV-mediated manipulation of MAP2K1 on hnRNP-K location and cellular survival. HIV-infected Ghost(3) cells (Figures 3D, S3C-S3D) and Mφ (Figures 3E, S3E-S3F) exposed to a MAP2K1 inhibitor revealed strong nuclear localisation of hnRNP-K as measured by immunofluorescence, with infected Mφ undergoing apoptosis in the presence of the inhibitor (Figures 3F, S3G). HIV-infected cells treated with an ERK2 inhibitor showed a similar hnRNP-K phenotype (data not shown) suggesting that the virus requires both MAP2K1 and ERK2 to secure the pro-survival pathway. ERK2 is known to interact with the pre-integration complex (Jacque et al., 1998; Bukong et al., 2010) as well as Tat soon after integration (Wolf et al., 2008). However, as apoptosis studies involving ERK2 are complicated by an unusually prolonged presence of activated (pro-survival) ERK2, specifically in the presence of the MAP2K1 inhibitor U0126 and Doxorubicin (Cagnol and Chambard, 2010), we did not monitor this host factor further. Instead, immunofluorescence staining of the Ser218/222 activation mark on MAP2K1 in HIV-infected Mφ showed distinct punctate cytoplasmic localisation of the protein, only in the presence of the inhibitor (Figures 3G, S3H). This was in line with prior art showing that aberrant sub-cellular localisation of MAP2K1/ERK pathway members is directly associated with cell death (Liu et al., 2008; Cagnol and Chambard, 2010). Taken together, these findings strongly supported our hypothesis that HIV-1 requires MAP2K1/ERK2 both for integration as well as to ensure cytoplasmic accumulation and sequestration of hnRNP-K, and therefore cellular survival. Given that MAP2K1 transcription is negatively regulated via the lincRNA-p21/hnRNP-K complex, by securing positive MAP2K1 signaling in concert with integration, HIV-1 is able to prevent nuclear entry of hnRNP-K, and thus prevent apoptosis.

Having identified the specific host protein manipulated by HIV-1 to ensure cytoplasmic hnRNP-K, we next sought to determine which viral protein could be mediating this effect. In contrast to HIV-1 expressing a native gp120 envelope protein, virus pseudotyped with vesicular stomatitis virus G (VSV-G) envelope protein was unable to exclude hnRNP-K from the nuclei of Mφ (Figure 3H) and Ghost(3) cells (Figure 3I). Supporting this observation, Ghost(3) cells transfected with gp120 envelope alone showed cytoplasmic hnRNP-K, while similarly treated cells transfected with HIV-1 Tat, did not (Figure 3I). Intriguingly, gp120 also affected HuR, such that VSV-G pseudotyped HIV-1 did not exclude HuR from the cytoplasm, and gp120 alone led to nuclear HuR (Figure S3I). Furthermore, VSV-G pseudotyped HIV-infected Mφ showed greater levels of apoptosis in the presence of a MAP2K1 inhibitor as compared to HIV-gp120 infected counterparts (Figure S3G). Together, these data strongly suggested that viral control of hnRNP-K’s cellular location requires gp120, and further demonstrated that HIV-1 gains control of hnRNP-K and HuR very early during infection.

### Restoring lncRNA/protein partner interactions in infected cells induces apoptosis

To confirm our MAP2K1/ERK2 findings, and conclusively demonstrate that cytoplasmic accumulation of hnRNP-K plays a central role in the mechanism HIV-1 uses to evade apoptosis, we interrogated the pathway by using drugs that regulate hnRNP-K activity in response to DNA damage and p53-mediated apoptosis. We designed an experiment by which we could restore hnRNP-K to the nucleus during HIV-1 infection and concurrently boost lincRNA-p21 levels. We reasoned that overcoming this block would help to re-establish hnRNP-K’s interaction with lincRNA-p21 and trigger cellular apoptosis in infected cells. DNA damage signaling culminates in hnRNP-K activation chiefly through inhibition of its ubiquitin-mediated degradation by the E3 ligase HDM2 (Moumen et al., 2005). Both p53 and hnRNP-K are thus activated in response to DNA damage and both are negatively regulated through the action of HDM2 (Enge et al., 2009). Nutlin3a is a small molecule inhibitor that activates p53 by binding to HDM2 (Vassilev et al., 2004). As Nutlin3a had previously been linked to activation of hnRNP-K (Moumen et al., 2005), we hypothesised that treatment with this inhibitor may allow hnRNP-K to move into the nucleus of HIV-infected cells (Figure 4A). Control Ghost(3) cells treated with Nutlin3a in the absence of any DNA damage showed no nuclear hnRNP-K revealing that even if the negative regulation mediated by HDM2 is released, DNA damage is still required to trigger nuclear localisation of hnRNP-K (Figure 4B upper panel). In contrast, Nutlin3a treatment following 24 hours of infection resulted in nuclear localisation of hnRNP-K in HIV-infected Ghost(3) cells only (Figure 4B lower panel). This result demonstrates that the addition of Nutlin3a after infection is sufficient to prevent the virus from excluding hnRNP-K from the nucleus of Ghost(3) cells.

**Figure 4.**
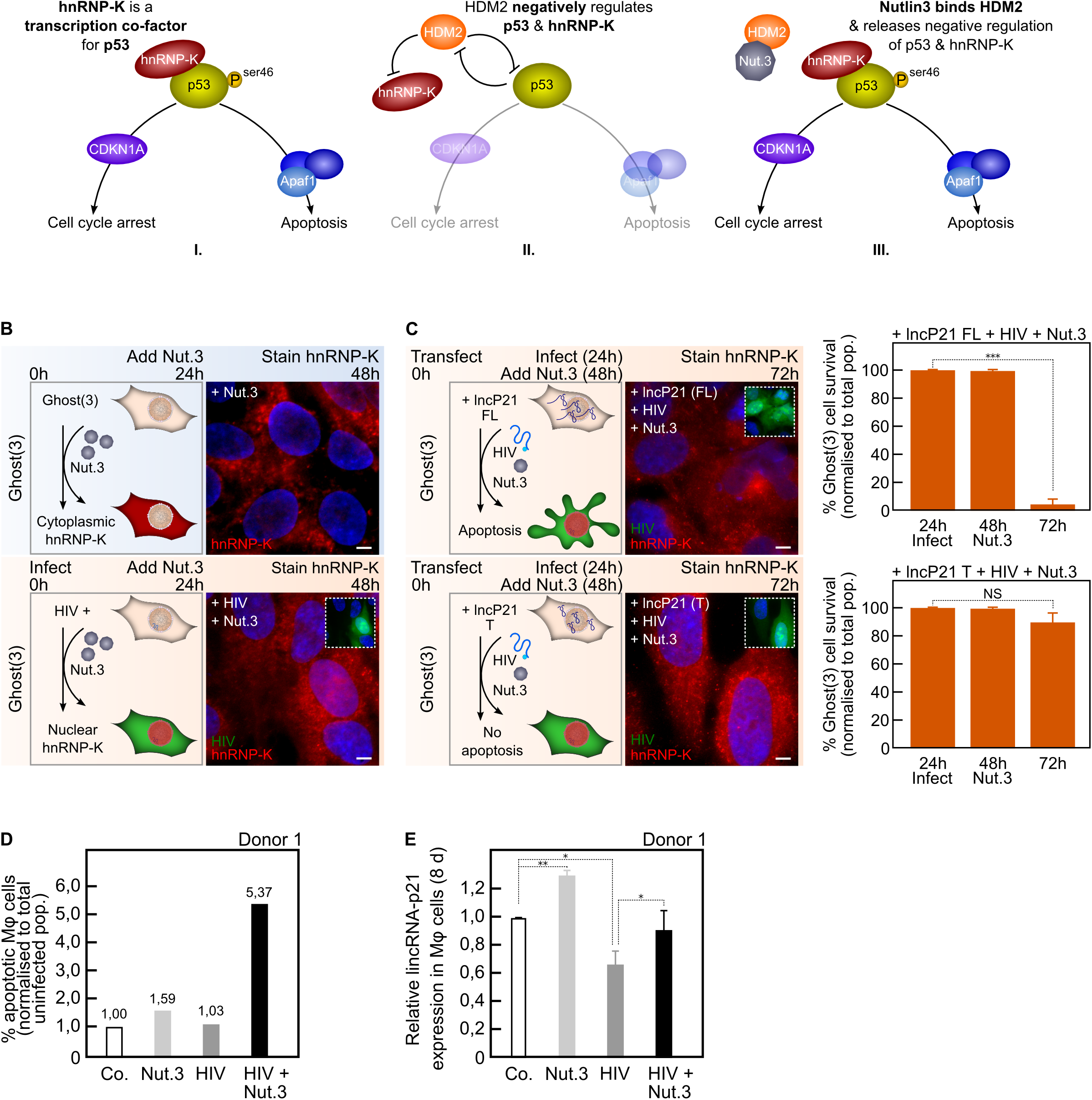
Nutlin3a confirms pivotal role of hnRNP-K in apoptosis evasion by HIV-1. (A) In response to DNA damage, both p53 and hnRNP-K are activated with cells undergoing apoptosis if p53ser46 is phosphorylated (I). In healthy cells, both p53 and hnRNP-K are negatively regulated by HDM2 (II). Nutlin3a (Nut.3) binds HDM2 thereby releasing its negative regulation of p53 and hnRNP-K (III). (B) Exposure of Ghost(3) cells to Nutlin3a 24 hours post-infection leads to nuclear localisation of hnRNP-K at 48 hours in infected (GFP-positive) cells only (+HIV+Nut.3a). Control cells show cytoplasmic hnRNP-K as expected in the absence of DNA damage. (C) Exogenous full-length lincRNA-p21 expression and Nutlin3a treatment lead to nuclear hnRNP-K and apoptosis in HIV-infected Ghost(3) cells (+lncP21-FL+HIV+Nut.3). Similarly treated cells transfected with truncated exogenous lincRNA-p21 show nuclear hnRNP-K but do not undergo apoptosis (+lncP21-T+HIV+Nut.3). (D) The percentage of apoptotic Mφ (normalised to total uninfected population) significantly increases only in HIV-infected Mφ exposed to Nutlin3a (HIV+Nut.3). Data shown for 1 of 3 donors. (E) LincRNA-p21 expression is significantly increased in HIV-infected Mφ exposed to Nutlin3a (HIV+Nut.3a) when compared to HIV-infection alone (+HIV) as detected by quantitative real-time RT-PCR analysis relative to the HPRT housekeeping gene and normalised to untreated cells (mean ± SE of 1 biological replicate in triplicate). Cells were counterstained with DAPI; scale bars = 10μM; see also supplemental figure 4.

Since HIV-1 infection lowers lincRNA-p21 levels by HuR-mediated degradation (Figures 2B-2C), overcoming the nuclear exclusion of hnRNP-K may be insufficient to induce apoptosis if lincRNA-p21 levels are not concomitantly up-regulated. Although Nutlin3a treatment could lead to nuclear localisation of hnRNP-K in the presence of HIV-1, we sought to combine this effect with elevated lincRNA-p21 expression, which was technically feasible in Ghost(3) cells. Importantly, we had previously observed that overexpressed lincRNA-p21 did not lead to nuclear hnRNP-K and apoptosis in HIV-infected Ghost(3) cells (Figures S4A-S4B). We thus transfected Ghost(3) cells for 24 hours with lincRNA-p21, infected for 24 hours, and followed with Nutlin3a treatment. Notably, only in the presence of full-length lincRNA-p21 did HIV-infected Ghost(3) cells treated with Nutlin3a demonstrate nuclear hnRNP-K and undergo apoptosis (Figure 4C upper panel). In contrast, treated cells that received a truncated version of lincRNA-p21, which is unable to bind hnRNP-K (Huarte et al., 2010), showed nuclear hnRNP-K but did not undergo apoptosis (Figure 4C lower panel). Furthermore, we verified that the combination of overexpressed lincRNA-p21 and Nutlin3a in the absence of HIV-1 did not lead to nuclear localisation of hnRNP-K and apoptosis in Ghost(3) cells (Figures S4B-S4C). Crucially, while exogenous overexpression of full-length or truncated lincRNA-p21 was not possible in Mφ, the addition of Nutlin3a in the presence of virus induced apoptosis (Figures 4D, S4D). Furthermore, this effect was particularly evident in Mφ infected with VSV-G pseudotyped HIV-1 (Figure S4D), in line with our previous data showing that gp120 plays a crucial role in the virus’ ability to evade hnRNP-K mediated apoptosis. Finally, lincRNA-p21 levels were concomitantly upregulated in HIV-infected Mφ treated with Nutlin3a (Figures 4E, S4E).

Overall, these results indicate that restoring hnRNP-K to the nucleus in the presence of elevated full-length lincRNA-p21, leads to apoptosis in HIV-infected cells. In addition, the observation that truncated lincRNA-p21 did not similarly result in apoptosis strongly supports the agency of lincRNA-p21 in this process, confirming the requirement of a fully functional binding complex comprised of lincRNA-p21 and hnRNP-K (Figure S4F) (Huarte et al., 2010). Furthermore, the enhanced apoptosis levels observed in the presence of VSV-G pseudotyped virus firmly support the role of gp120 in this mechanism. In sum, these results highlight the pivotal role that a host long noncoding RNA plays in HIV-1 apoptosis evasion mechanisms.

## DISCUSSION

Our findings elucidate the deliberate and focused attack HIV-1 aims at lincRNA-p21 to mask IN-induced DSBs and evade cellular apoptosis (Figure 1). The virus does so very early in its infectious cycle by hijacking cellular HuR to degrade lincRNA-p21, and cripples its association with the nuclear protein binding partner hnRNP-K by spatially separating the two molecules (Figure 2). HIV-1 ensures cytoplasmic accumulation of hnRNP-K via the previously demonstrated but until now poorly understood action of MAP2K1 and ERK2 (Figure 3). Appropriate nuclear interaction between lincRNA-p21 and hnRNP-K can be restored through the inhibition of MAP2K1 or ERK2 (Figure 3), or via the combined action of Nutlin3a and enhanced expression of lincRNA-p21 (Figure 4). We propose a model (Figure 5) in which HIV-1 secures the pro-survival pathway by ensuring activated MAP2K1 continues to phosphorylate ERK2 thereby leading to cytoplasmic accumulation of hnRNP-K as well as phosphorylation of HDM2. HIV-1 had been previously reported to use this pathway to facilitate integration through the viral Env protein gp120, though the significance of these interactions was not fully appreciated until now.

**Figure 5.**
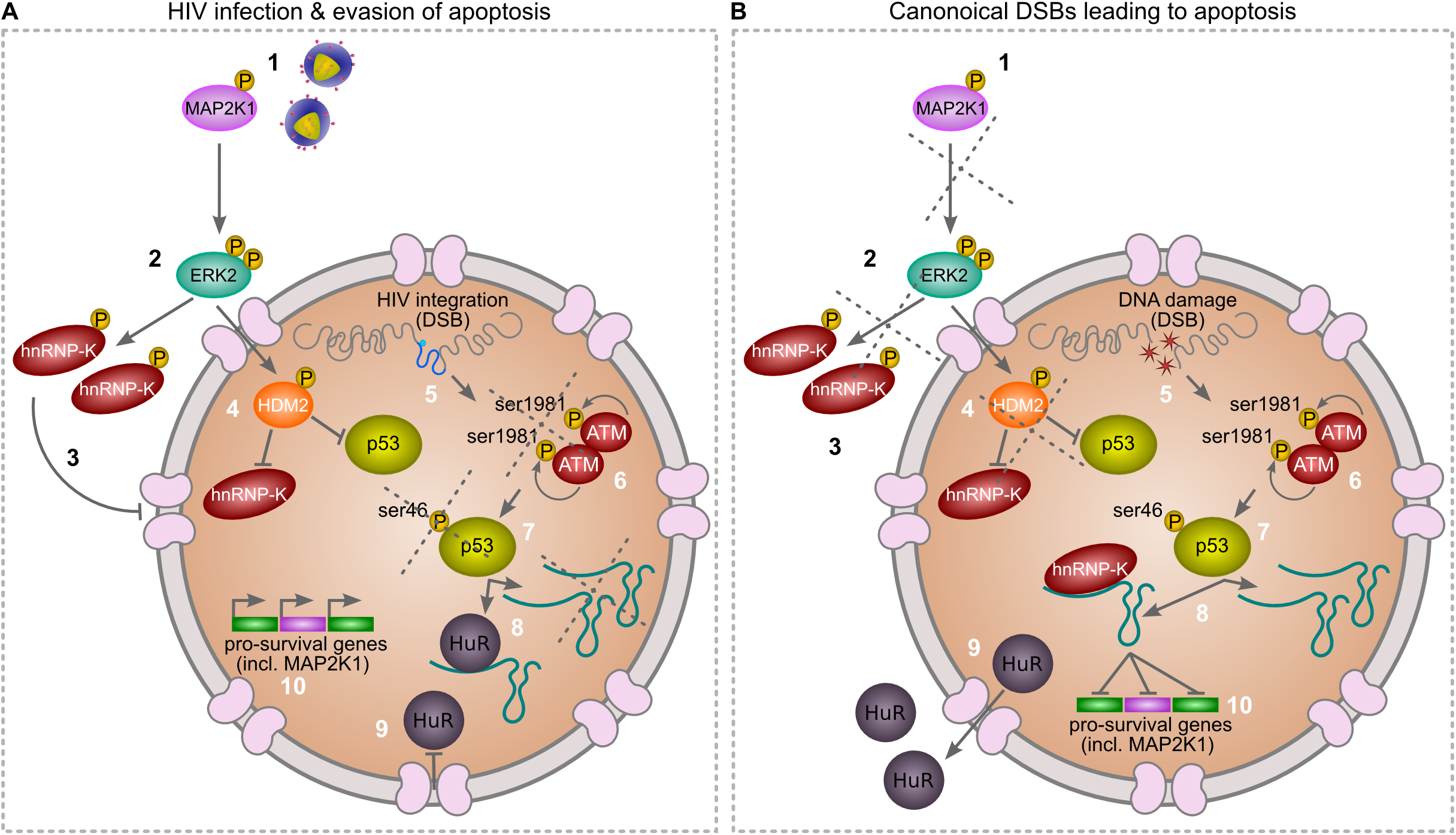
Hypothetical model of HIV-mediated manipulation of lincRNA-p21 to evade cellular apoptosis. During canonical DNA damage, the MAP2K1/ERK2 pathway is inactivated. However, during infection, HIV-1 ensures activated MAP2K1 (1) continues to phosphorylate ERK2 (2) thereby leading to cytoplasmic accumulation of hnRNP-K (3) as well as phosphorylation of HDM2 (4). The latter action of ERK2 ensures ubiquitin-mediated degradation of nuclear hnRNP-K as well as p53. When compared to canonical DNA damage, both HIV-1 integration and Doxorubicin induce DSBs (5). However, in HIV-infected cells this does not lead to autophosphorylation and activation of ATM at serine1981 (6) or subsequent phosphorylation of p53 at serine46 (7). The p53pSer46 apoptosis mark leads to increased lincRNA-p21 transcription and association with hnRNP-K (8), as well as translocation of HuR to the cytoplasm (9) of canonically DNA damaged cells only. Nuclear lincRNA-p21/hnRNP-K complexes lead to suppression of pro-survival genes in canonically DNA damaged cells only (10). As HIV-1 is able to alter the location of hnRNP-K (3) and HuR (9), lincRNA-p21 expression is low and pro-survival genes, including MAP2K1, are not repressed in infected cells (10), thus the virus is able to evade apoptosis.

Our data strongly suggest that HIV-1 can secure the pro-survival pathway in non-CD4+ T cells very early in the viral life cycle, a pathway that is unavailable to HIV-1 in CD4+ T cells due to the well-reported absence of ERK2 expression following differentiation (Fischer et al., 2005; Chang et al., 2012). In support of this, lincRNA-p21 expression was indeed increased in HIV-infected CD4+ T cells (data not shown) and Jurkat cells (Imam et al., 2015). The early stage at which HIV-1 secures that pathway is supported by the documented interaction of HIV-1 IN and Tat with ERK2 (Jacque et al., 1998; Bukong et al., 2010; Wolf et al., 2008) as well as our own gp120 data (Figure 3), although the use of various gp120 mutants would probably illuminate this more robustly. Furthermore, our data show that HIV-1 integration does not lead to complete p53 activation, specifically phosphorylation of Ser46. As a consequence, transcription of pro-apoptotic lincRNA-p21 is decreased, and the lincRNA-p21/hnRNP-K feedback loop that suppresses pro-survival MAP2K1, is inactive.

Our data provide a glimpse of how pathogens with limited coding capacity can significantly restructure major cellular pathways using host lncRNAs. This is even more striking in the context of cell type-specific transcription: activated CD4+ T cells express low levels of MAP2K1/ERK2, and are highly sensitive to HIV-induced apoptosis. In contrast, MAP2K1/ERK2 expression remains high in macrophages (Chang et al., 2012), and these cells robustly resist viral-induced apoptosis. Thus our data suggest that the lack of ERK2 expression in CD4+ T cells may be a key reason why HIV-1 is not able to evade apoptosis in these cells, while doing so in macrophages. MAP2K1 and lincRNA-p21 are at the nexus of these pathways and their manipulation by HIV-1 adds a new layer of complexity to host-pathogen interactions.

Our findings also highlight that three discrete but tightly coordinated events are required for apoptosis to occur. Firstly, and already well-established, DNA damage such as DSBs must lead to complete activation of p53 such that downstream transcription of associated targets is initiated. Secondly, lincRNA-p21 expression levels must be enhanced. This is connected to p53 activation and HuR-mediated degradation, but can be bypassed through exogenous overexpression. Third and lastly, hnRNP-K must translocate to the nucleus to ensure its association with lincRNA-p21 and subsequent suppression of pro-survival genes such as MAP2K1. An absence of any one of these events is sufficient for the evasion of apoptosis as summarised by our findings (Figure S4F).

While it is not unusual for pathogens to manipulate host signaling pathways, our data illuminate a unique strategy whereby HIV-1 is able to control the cellular response to DNA damage via a long noncoding RNA. It has been observed that Adenovirus oncoproteins are able to inactivate the DNA repair MRN complex at viral replication centers, masking host genome instabilities that are instigated by this generally non-integrative DNA virus (Stracker et al., 2002; Shah and O’Shea, 2015). Given that viruses contribute to 20% of cancers worldwide (de Martel et al., 2012), it is important to understand how genomic instabilities are propagated following challenge, as well as how apoptosis is evaded. While HIV-1 itself does not cause cancer, our observations that HIV-mediated resistance to apoptosis occurs through manipulation of a host long noncoding RNA and its protein binding partners, reveals a novel mechanism whereby genomic integrity can be severely challenged yet cells survive. In addition, we suspect that gp120 may be initiating a general DNA damage suppression phenotype via lincRNA-p21, possibly to aid macrophage-mediated dissemination throughout the host. Lastly, given the uncontested connection between infection and cancer (de Martel et al., 2012), our data provides a novel perspective on how pathogens may increase DNA damage while simultaneously blocking apoptosis.

## EXPERIMENTAL PROCEDURES

### Cell culture

Ghost(3) cells (AIDS Research and Reference Reagent Program, Division of AIDS, NIAID, NIH) were cultured in DMEM (Invitrogen) supplemented with 10% heat-inactivated fetal bovine serum (FBS, Biochrom), 0.2mM GlutaMAX™ (Life Tech), 500μg/mL G418 (Sigma), 100μg/mL Hygromycin (Sigma) and 1μg/mL Puromycin (Sigma). HEK293T cells (AIDS Research and Reference Reagent Program, Division of AIDS, NIAID, NIH) were cultured in DMEM (Invitrogen) supplemented with 10% heat-inactivated FBS (Biochrom). PBMCs from healthy anonymous donors were isolated from buffy coats processed by the Western Province Blood Transfusion Service (University of Cape Town HREC ref 317/2016). PBMCs were separated by density centrifugation over Lymphoprep (Alere). Monocytes were isolated were isolated using Ficoll (GE Healthcare) followed by CD14+ positive selection (Miltenyi microbeads), cultured and differentiated into macrophages for 7 days in RPMI (Invitrogen) supplemented with 10% heat-inactivated Human AB serum (IPLA), sodium pyruvate (Invitrogen), L-Glutamine (Invitrogen) and 20ng/mL M-CSF (Miltenyi).

### Viral plasmids, virus stocks, infections and drugs

Viral stocks were generated by co-transfecting HEK293T cells with either HIV-1 clones BaL.01 and pSG3Δenv, or NL4-3 and AD8env (AIDS Research and Reference Reagent Program, Division of AIDS, NIAID, NIH) using Fugene6 (Roche). Supernatants were collected 48 hours post-transfection, supplemented with FBS to a final concentration of 20% and stored in aliquots at -80°C. HEK293T cell viral stocks were used to infect Ghost(3) cells at an MOI = 1.0 for up to 72 hours. For human macrophage infections, HIV-1 strain BaL from National Institute for Biological Standards and Controls was propagated in PBMCs in RPMI-supplemented with 50 mM glutamine,20% FBS, 100 IU/mL penicillin, 100 μg/mL streptomycin, and 20 IU/mL IL-2 (Sigma) (34). Supernatants were filtered through a 0.22-μM PVDF membrane (Millipore) and purified by ultracentrifugation through a 20% sucrose buffer and resuspended in RPMI medium with 5% human AB serum. Mφ were infected to a final rate of ~ 4,5% for up to 9 days. Virus input was washed out 24 hours post-infection of Ghost(3) cells and three days post-infection for Mφ. Where indicated, cells were treated with 10μM of either Doxorubicin (Sigma-Aldrich), Nutlin3a (Sigma-Aldrich), MAP2K1 inhibitor (U-0126) or ERK2 inhibitor (FR180204) for up to 48 hours.

### Cloning and transfections

Ghost(3) cells were transfected (for 48 hours prior to infection) with 25nM (final concentration) of ON-TARGETplus human HuR/ELAV1 siRNA SMARTpool (5’ GAC AAA AUC UUA CAG GUU U 3’, 5’ GAC AUG UUC UCU CGG UUU G 3’, 5’ ACA AAU AAC UCG CUC AUG C 3’, 5’ GCU CAG AGG UGA UCA AAG A 3’; ThermoScientific) using RNAiMax (Invitrogen). Mouse full-length lincRNA-p21 (3073 bp) and truncated lincRNA-p21 (1889 bp) sequences (7) were synthesised (GeneArt, Life Technologies) and sub-cloned via 5’ SacI and 3’ EcoRI into pCi-Neo (Promega). Ghost(3) cells were transfected for 21 hours (prior to infection) with either construct using Lipofectamine2000 (Invitrogen).

### Immunofluorescence

For each experiment, cells were infected or treated with drugs on coverslips, fixed for 10 mins in fresh 4% paraformaldehyde at room temperature, then washed 3 times in PBS and permeabilised for 10 mins in ice-cold methanol at -20°C. Coverslips were washed once in PBS and incubated in blocking buffer (5% goat serum, 0.3% Triton-X100 in PBS) for 60 mins at room temperature. Cells were incubated in primary antibody solution (1% BSA, 0.3% Triton X-100 in PBS) overnight at 4°C using the following antibodies rabbit polyclonal anti-phospho-histone H2A.X Ser139 (Cell Signaling), rabbit monoclonal anti-phospho-ATM Ser1981 (Cell Signaling), rabbit polyclonal anti-phospho-p53 Ser46 (Cell Signaling), mouse monoclonal anti-HIV-1 p24 (AIDS Research and Reference Reagent Program, Division of AIDS, NIAID, NIH), mouse monoclonal anti-HuR 3A2 (Santa Cruz Biotechnology), goat polyclonal anti-hnRNP-K P-20 (Santa Cruz Biotechnology), and rabbit polyclonal anti-phospho-MAP2K1/MAP2K2 Ser218/Ser222 (Elabscience Biotechnology). Coverslips were washed 3 times (5 mins each on an orbital shaker) with wash buffer (0.05% Tween-20 in PBS), followed by incubation with secondary antibodies conjugated to either Atto-550 or Atto-565 or Atto-647 for 60 mins at room temperature. Coverslips were washed 3 times (5 mins each on an orbital shaker) with wash buffer (0.05% Tween-20 in PBS). Coverslips were incubated in equilibration buffer (0.4% glucose, 2X SSC) for 5 mins and counterstained with 1mg/ml DAPI (4’,6-diamidino-2-phenylindole; Life Technologies). Coverslips were mounted in glox buffer (3.7mg/ml glucose oxidase, 1U catalase) and imaged. A range of 60 to 600 cells per treatment were imaged as described below.

### Quantitative RT-PCR

For each experiment, cells were infected or treated with drugs over time and total RNA was extracted from cells using TRIzol^®^ (Life Technologies) and Direct-zol™ (Zymo Research), treated with RQ1 RNase-free DNase (Promega) and reverse transcribed using SuperScript III Reverse Transcriptase (Life Technologies). Quantitative RT-PCR was performed using Sso Fast EvaGreen™ supermix (BioRad) on a Bio-Rad CFX96 real-time PCR detection system. PCR primer sets included: p53 (forward 5’ TGTGACTTGCACGTACTCCC 3’, reverse 5’ ACCATCGCTATCTGAGCAGC 3’), CDKN1A/p21 (forward 5’ AGTCAGTTCCTTGTGGAGCC 3’, reverse 5’ GACATGGCGCCTCCTCTG 3’), HIV-1 Gag (5’ ACTCTAAGAGCCGAGCAAGC 3’, reverse 5’ TGTAGCTGCTGGTCCCAATG 3’), lincRNA-p21 (forward 5’ GGGTGGCTCACTCTTCTGGC 3’, reverse 5’ TGGCCTTGCCCGGGCTTGTC 3’), HuR (forward 5’ AGAGCGATCAACACGCTGAA 3’, reverse 5’ TAAACGCAACCCCTCTGG AC 3’), MAP2K1 (forward 5’ ATGGATGGAGGTTCTCTGGA 3’, reverse 5’ TTTCTGGCGACATGTAGGACC 3’) and HPRT (forward 5’ GCAGCCCTGGCGTCGTGATTA 3’, reverse 5’ CGTGGGGTCCTTTTCACCAGCA 3’).

### Apoptosis assay

Apoptosis was measured in Ghost(3) cells 30 hours post-infection or drug treatment using the NucViewTM 488 Caspase-3 Assay Kit (Biotium) in a 96 well format. Data was analysed in MATLAB using a probabilistic region method to measure cell attachment and expressed as percentage cell survival. Notably, as the kit was only available with a 488nM dye, cells were analysed at the 30 hour time-point to minimise GFP input from the integrated Tat-driven reporter. An average of 3000 cells per condition were analysed with the exception of cells treated with Doxorubicin first followed by HIV-1 infection (Figure 1A) and cells transfected with lincRNA-p21 overexpression constructs followed by Doxorubicin treatment (Figure S2H). These conditions (⌘) yielded too few attached cells (<20) at 30 hours for similar analysis. Apoptosis was measured in Mφs 9 days post-infection and/or 2 days after drug treatment using the fixable viability dye Aqua Live/Dead (Thermo Fisher Scientific) in combination with the FLICA 660 caspase 3/7 dye (Thermo Fisher Scientific) after detachment using StemPro^®^ Accutase^®^ (Thermo Fisher Scientific). Cells were then treated with cytofix/cytoperm and perm/wash solutions (BD Biosciences) and stained with FITC-conjugated anti-p24 antibody. A minimum of 10^4^ events were acquired per treatment on a BD LSRForstessa™ flow cytometer.

### RNA pulldown and mass spectrometry

RNA pulldown and mass spectrometry was performed as previously described (Huarte et al., 2010) using Ghost(3) cell lysate and 50pmol of biotinylated probes targeted to lincRNA-p21.

### Imaging and analysis

Cells were imaged on one of two microscopes: 1) a customised Nikon Ti Eclipse widefield fluorescent microscope using a 100x 1.49 N.A. Nikon Apochromat TIRF oil immersion objective, with mercury lamp illumination through the appropriate Semrock razor sharp filter sets at low camera gain in each of the fluorescent channels using an Andor iXion 897 EMCCD camera cooled to -80°C, and controlled using μmanager open source microscope management software (NIH and UCSF, USA). A 30ms exposure time was used for DAPI. Exposure times ranged from 200 to 500ms for other dyes. Each field of view was captured as a series of images acquired on multiple focal planes through the samples, across a range of 2-10mm in the axial plane. A 0.2mm piezo step-size was used for z-stacks. 2) An Andor Technology (Belfast, Northern Ireland) integrated Yokogawa CSU-W1 spinning disk confocal system and a Metamorph controlled Nikon TiE motorized microscope with a 60x, 1.4 NA phase oil immersion objective. Excitation sources were 405, 488 and 561 laser lines and emissions were detected through Semrock Brightline 465, 525 and 607 nm filters. Images were captured using an 888 EMCCD camera (Andor) and iQ2 software. Eight fields of view were captured per well (two wells per condition, from 3 donors), eight Z-sections per FOV with one phase and three fluorescent images. Signal intensities of all images were measured using Fiji (Schindelin et al., 2012). The contrast of images shown was adjusted to fit a 16 bit gray-scale.

## AUTHOR CONTRIBUTIONS

S.B. and M.M.M. designed the experiments. S.B., J.N., J.S., M.B, S.P. and A.K.C conducted the experiments. S.B. and M.M.M. analysed the data. F.B. supervised S.P. S.B. and M.M.M. wrote the manuscript. M.M.M. supervised the overall study.

## ACKNOWLEDGEMENTS

We thank all members of the Gene Expression and Biophysics Laboratory. We thank A. Savulescu and S. Stoychev and N. Peton for technical assistance and for reagents. We thank S. Fanucchi, M. S. Weinberg for comments on the manuscript. S.B is a recipient of a Sydney Brenner Postdoctoral Fellowship. J.S. is a recipient of a NRF/PDP Postdoctoral Fellowship. A.S. is a recipient of the Human Frontiers Science Organization Career Development Award. This research is supported by grant PG V2KYP07 from the Council for Industrial and Scientific Research (CSIR, South Africa) and by a grant from the Emerging Research Area Program & the Centre of Competence Program of The Department of Science and Technology (DST, South Africa) and a grant from the Medical Research Council (South Africa) all to M.M.M.

## FIGURE LEGEND

**Supplemental figure 1. HIV-1 masks DNA damage, protects against additional lethal DNA damage, and prevents lincRNA-p21 upregulation**

(A) HIV-1 integration occurs approximately 16 hours post-infection of Ghost(3) reporter cells and Tat-mediated activation of an integrated LTR-driven GFP reporter can be detected approximately 48 hours post-infection.

(B) Ghost(3) cells support HIV-1 replication over time as detected by quantitative real-time RT-PCR analysis of HIV-1 Gag expression relative to the HPRT housekeeping gene and normalised to uninfected cells (mean ± SE of 3 biological replicates in triplicate).

(C) HIV-1 infection of Ghost(3) cells induces DSBs over 48 hours as detected by H2A.XpSer139 immunofluorescence staining but does not lead to caspase 3-mediated apoptosis (displayed as percentage cell survival; n = 3000). Doxorubicin treatment (Doxo.) over the same time course yields extensive H2A.XpSer139 staining followed by apoptosis.

(D) HIV-mediated DSBs require integration as addition of Raltegravir (Ralt.), an integrase inhibitor, prevents DSBs and H2A.XpSer139 staining in HIV-infected Ghost(3) cells over 48 hours.

(E) Raltegravir (Ralt.) prevents HIV-1 infection of Ghost(3) cells as detected by quantitative real-time RT-PCR analysis of HIV-1 Gag expression relative to the HPRT housekeeping gene and normalised to uninfected cells (mean ± SE of 3 biological replicates in triplicate).

(F) Nuclear inactive p53 monomers are not phosphorylated at serine residue 46 (specific apoptotic mark) in response to HIV-1 infection of Mφ as measured by immunofluorescence staining (p53pSer46). Nuclear activated p53 dimers are detected in Doxorubicin-treated cells. Cells were counterstained with DAPI; scale bars = 10μM; two-tailed paired Student T-test, ***p < 0.001, **p < 0.01, *p < 0.05, NS (not significant).

**Supplemental figure 2. HIV-1 manipulates lincRNA-p21’s protein binding partners**

(A) Untreated Ghost(3) cells show nuclear HuR over a 48 hour time course by immunofluorescence staining.

(B) HuR expression is significantly decreased as measured by quantitative real-time RT-PCR analysis following 48 hours of exposure to siHuR in Ghost(3) cells (mean ± SE of 3 biological replicates in triplicate).

(C) LincRNA-p21 expression increases in the absence of HuR in untreated and HIV-infected Ghost(3) cells as measured over time by quantitative real-time RT-PCR analysis relative to the HPRT housekeeping gene (mean ± SE of 3 biological replicates in triplicate).

(D) siHuR-treated Ghost(3) cells support HIV-1 replication to the same extent as untreated cells, as indicated by GFP expression. Scale bar = 5μM.

(E) siHuR-treated Ghost(3) cells support HIV-1 replication as measured by quantitative real-time RT-PCR analysis of HIV-1 Gag relative to the HPRT housekeeping gene (mean ± SE of 3 biological replicates in triplicate).

(F) Exogenous full-length lincRNA-p21 expression is significantly decreased in the presence of HIV-1 as measured over time in Ghost(3) cells by quantitative real-time RT-PCR analysis relative to HPRT housekeeping gene (mean ± SE of 3 biological replicates in triplicate).

(G) Exogenous full-length lincRNA-p21 treated Ghost(3) cells support HIV-1 replication as measured by quantitative real-time RT-PCR analysis of HIV-1 Gag relative to the HPRT housekeeping gene (mean ± SE of 3 biological replicates in triplicate).

(H) Exogenous full-length lincRNA-p21 expression (FL) followed by Doxorubicin treatment leads to apoptosis in Ghost(3) cells. No other treatments lead to significant apoptosis. ⌘ Too few attached cells (< 20) were present for statistical analysis.

(I) Schematic representation of RNA pulldown and mass spectrometry experiments used to identify protein binding partners of lincRNA-p21 in the presence of HIV-1. Biotinylated probes targeted to lincRNA-p21 were incubated with cellular extracts, targeted using streptavidin beads, washed, resolved on a polyacrylamide gel and identified by mass spectrometry.

(J) In uninfected Ghost(3) cells, lincRNA-p21 associated with a unique set of proteins (red circle). Similarly, in the presence of HIV-1, lincRNA-p21 associated with a different unique set of proteins (green circle). Another subset of proteins associated with lincRNA-p21 both in the presence and absence of HIV-1 but at different levels. In the presence of HIV-1, hnRNP-K (red) associated less with lincRNA-p21. In the presence of HIV-1, HuR, XRCC6 and PRKDC (green) associated more with lincRNA-p21.

Cells were counterstained with DAPI; scale bars = 10μM (Figure S2D, 5μM); two-tailed paired Student T-test, ***p < 0.001, **p < 0.01, *p < 0.05, NS (not significant).

**Supplemental figure 3. HIV-1 requires gp120 Env and MAP2K1/ERK2 to ensure hnRNP-K’s cytoplasmic localisation**

(A) Quantitative real-time RT-PCR analysis of MAP2K1 expression relative to HPRT housekeeping gene in MAP2K1 inhibitor-treated (MPK inh.), or ERK2 inhibitor-treated (ERK inh.), or Doxorubicin-treated (+Doxo.) Ghost(3) cells normalised to untreated cells (mean ± SE of 3 biological replicates in triplicate).

(B) Quantitative real-time RT-PCR analysis of lincRNA-p21 expression relative to HPRT housekeeping gene in MAP2K1 inhibitor-treated (MPK inh.) or ERK2 inhibitor-treated (ERK inh.) Ghost(3) cells normalised to untreated cells (mean ± SE of 3 biological replicates in triplicate).

(C) Inhibition of MAP2K1 allows for nuclear localisation of hnRNP-K as measured by immunofluorescence staining, but no apoptosis occurs in treated Ghost(3) cells.

(D) Quantification of nuclear localised hnRNP-K in infected (HIV), treated (MPK inh.) or infected and treated (HIV+MPK inh.) Ghost(3) cells shown as mean relative fluorescence intensity (RFI) ± SE of 3 biological replicates.

(E) Inhibition of MAP2K1 allows for nuclear localisation of hnRNP-K as measured by immunofluorescence staining, but no apoptosis occurs in treated Mφ.

(F) Quantification of nuclear localised hnRNP-K in infected (HIV), treated (MPK inh.) or infected and treated (HIV+MPK inh.) Mφ, shown as mean relative fluorescence intensity (RFI) ± SE of 3 donors.

(G) The percentage of apoptotic Mφ (normalised to total uninfected population) increases significantly in cells infected with VSV-G pseudotyped HIV-1 and exposed to a MAP2K1 inhibitor (HIV VSV-G +MPK inh.). Mean ± SE of 3 donors.

(H) Immunofluorescence staining of the Ser218/222 activation mark on MAP2K1 in Mφ or those treated with an inhibitor (+MPK inh.) after 8 days.

(I) HIV-1 clone BaL.01 packaged with gp120 Env (+HIV gp120) sequesters HuR in the nucleus of infected Ghost(3) cells as shown by immunofluorescence staining, while VSV-G pseudotyped HIV-1 does not. Cells transfected with a plasmid expressing only HIV-1 gp120 (+gp120) show nuclear HuR while control cells transfected with a plasmid expressing only HIV-1 Tat (+Tat) show cytoplasmic and nuclear HuR.

Cells were counterstained with DAPI; scale bars = 10μM. two-tailed paired Student T-test, ***p < 0.001, **p < 0.01, *p < 0.05, NS (not significant).

**Supplemental figure 4. Nutlin3a confirms pivotal role of hnRNP-K in apoptosis evasion by HIV-1**

(A) Exogenous lincRNA-p21 (full-length or truncated) expression alone does not lead to nuclear hnRNP-K in the presence of HIV-1 in Ghost(3) cells.

(B) Exogenous lincRNA-p21 (full-length or truncated) expression and Nutlin3a treatment do not lead to apoptosis in Ghost(3) cells.

(C) Exogenous lincRNA-p21 (full-length or truncated) expression and Nutlin3a treatment do not lead to nuclear hnRNP-K in Ghost(3) cells.

(D) The percentage of apoptotic Mφ (normalised to total uninfected population) significantly increases in Mφ infected with VSV-G pseudotyped HIV-1 and exposed to Nutlin3a (HIV VSV-G+Nut.3). Mean ± SE of 3 donors.

(E) LincRNA-p21 expression is significantly increased in Mφ infected with either gp120 or VSV-G pseudotyped HIV-1 and exposed to Nutlin3a, as detected by quantitative real-time RT-PCR analysis relative to the HPRT housekeeping gene and normalised to untreated cells (mean ± SE of 2 biological replicates in triplicate).

(F) Summary of experimental conditions showing that only a combination of DNA damage, enhanced lincRNA-p21 expression and nuclear hnRNP-K leads to apoptosis. An absence of any one of these events, or exogenous expression of truncated lincRNA-p21 (which is unable to bind hnRNP-K), is sufficient to prevent apoptosis.

Cells were counterstained with DAPI; scale bars = 10μM. two-tailed paired Student T-test, ***p < 0.001, **p < 0.01, *p < 0.05, NS (not significant).

